# Identification of SIRT4 as a novel paralog-specific interactor and candidate suppressor of C-RAF kinase in MAPK signaling

**DOI:** 10.1101/2023.11.23.568463

**Authors:** Mehrnaz Mehrabipour, Radovan Dvorsky, Saeideh Nakhaei-Rad, Alexander Lang, Patrick Verhülsdonk, Mohammad Reza Ahmadian, Roland P. Piekorz

**Author notes:** Department of Cardiology, Pulmonology, and Vascular Medicine, Medical Faculty and University Hospital Düsseldorf, Heinrich Heine University, 40225 Düsseldorf, Germany (Alexander Lang). Equal contribution (Mohammad Reza Ahmadian, Roland P. Piekorz).

## Abstract

Cellular responses leading to development, proliferation, and differentiation rely on RAF/MEK/ERK signaling that integrates and amplifies signals from various stimuli to cellular downstream responses. The clinical significance of C-RAF activation has been reported in many types of tumor cell proliferation and developmental disorders, which requires the discovery of potential C-RAF protein regulators. Here, we identify a novel and specific protein interaction between C-RAF, among the RAF kinase paralogs, and SIRT4 among the mitochondrial sirtuin family members SIRT3, SIRT4, and SIRT5. Structurally, C-RAF binds to SIRT4 through the N-terminal cysteine-rich domain (CRD; a.a. 136-187), and on the other side, SIRT4 requires predominantly the C-terminus (a.a. 255-314) for full interaction with C-RAF. Interestingly, SIRT4 interacts specifically with C-RAF in a pre-signaling inactive (serine 259 phosphorylated) state. Consistent with this finding, ectopic expression of SIRT4 in HEK293 cells results in upregulation of pS259-C-RAF levels and concomitant reduction of MAPK signaling as evidenced by strongly decreased phospho-ERK signals. Thus, our findings propose another extra-mitochondrial role of SIRT4 and suggest that SIRT4 functions as a cytosolic tumor suppressor of C-RAF-MAPK signaling, besides its known metabolic tumor suppressor role towards glutamate dehydrogenase and glutamine levels in mitochondria.

## Introduction

C-RAF (often also called RAF1) belongs to the RAF kinases family (A-RAF, B-RAF, and C-RAF) which transfer proliferative and growth signals to downstream activation of MEK/ERK kinases. These RAF paralogs share several structural properties (1,2), yet they differ in terms of activity levels and functional roles (3). Among them, C-RAF exhibits moderate activity, less than B-RAF, but more than A-RAF, and is associated with cancer and developmental disorders (4–7). There are three conserved regions (CR) within RAF proteins that are important for their respective regulatory functions (CR1 and CR2) and kinase activity (CR3) (1). CR1 contains a RAS binding domain (RBD), mediating RAS interaction, and a cysteine-rich domain (CRD), which mediates membrane binding and enhances RAS/RBD affinity at the membrane (8–10). CR2 is enriched by several Ser/Thr residues, including serine 259 (S259) which is an important site for inhibitory phosphorylation and 14-3-3 binding that regulates RAF kinase activation (11). When phosphorylated by upstream kinases such as AKT, PKA, or LATS1, CR2 acts as an inhibitory domain that keeps C-RAF in an inactive state (12–14). Dephosphorylation of CR2 by protein phosphatases, such as PP2A or PP1, relieves this autoinhibition on the kinase domain and activates C-RAF (15). CR3 functions as a catalytic C-terminal region, constituting a putative phosphorylation segment for kinase activation (16). Thus, C-RAF cycles between a close inactive and an open active conformation which is regulated by different phosphorylation and dephosphorylation events (17). Overall, phosphorylation, feedback/autoinhibition, and protein-protein interaction occur in C-RAF regulation in response to signaling events (12,17–21). In particular, RAS and 14-3-3 binding are major regulatory events of RAF activation, membrane recruitment, and stabilization (8,22–24). Addressing the molecular control of C-RAF by interacting regulators and the underlying molecular and structural mechanisms is still necessary for understanding the complex landscape of Mitogen-Activated Protein Kinase (MAPK) network signaling. Several proteins that bind and regulate C-RAF have been identified, including RKIP (RAF1 kinase inhibitor protein) which functions as an anti-metastatic tumor suppressor and is downregulated in various cancers (25–28). RKIP binds to the N-terminal region of C-RAF and thereby inhibits C-RAF mediated phosphorylation and activation of MEK1/2 (29).

The family of human sirtuins comprises seven members, of which SIRT3, SIRT4, and SIRT5, function as *bona fide* metabolic regulators in mitochondria (30). In particular, SIRT4 inhibits, as a tumor suppressor, the metabolic gate-keepers pyruvate dehydrogenase (PDH) and glutamate dehydrogenase (GDH) (31,32), with particular significance for the regulation of glutamine metabolism in tumor cells. Recent findings uncovered novel extra-mitochondrial roles of SIRT4 in microtubule dynamics and regulation of mitotic cell cycle progression, WNT/β-Catenin and Hippo signaling, and SNARE complex formation required for autophagosome-lysosome fusion (33–36). Interestingly, proteomic analysis of the SIRT4 interactome identified C-RAF as a potential binding partner of SIRT4, indicating a novel role of SIRT4 in the regulation of the RAF-MAPK signaling pathway (33). Consistent with this idea, recent studies have demonstrated that (i) the tumor suppressor SIRT4 is downregulated in most human solid tumor types and cell lines (37–39), and (ii) ectopic expression of SIRT4 downregulates pERK1/2 levels and hence inhibits MAPK signaling and cell proliferation (37–42). Considering these interrelated findings, in this study we investigated the molecular and functional interaction between the proto-oncogene C-RAF and the tumor suppressor SIRT4 in the context of MAPK signaling inhibition.

## Material and methods

### Plasmid constructs

The N-terminal RBD-CRD, RBD, and CRD domains of RAF kinases were cloned into the pGEX-4T1 vector (BioCat GmbH). Upon transformation into *Escherichia coli* lysates containing GST-tagged proteins were prepared as previously described (43). The SIRT4 deletion mutants SIRT4(Δ69-98) (Δ1), SIRT4(Δ165-229) (Δ2), and SIRT4(Δ255-314) (Δ3) were generated by PCR-mediated mutagenesis and cloned into pcDNA-3.1 for eukaryotic expression as C-terminal eGFP fusion proteins. The expression construct for N-terminally Flag-tagged C-RAF was kindly provided by Dr. Motta (Genetics and Rare Diseases Research Division, Rome).

### Cell culture and generation of stable cell lines

HEK293 cells were maintained in Dulbecco’s modified Eagle’s serum (DMEM) (Thermo Fisher Scientific, Oberhausen, Germany) supplemented with 10% FBS (Gibco) and 1% Penicillin/Streptomycin (Genaxxon, Ulm, Germany). HEK293 cell lines stably expressing eGFP or C-terminally tagged SIRT4-eGFP or SIRT4(ΔN28)-eGFP have been previously described (44). In addition, HEK293 cell lines expressing Flag M2 as control or C-terminally Flag M2-tagged SIRT3, SIRT4, or SIRT5 proteins have been described (33). HEK293 cell lines stably expressing SIRT4(Δ69-98)-eGFP (Δ1), SIRT4(Δ165-229)-eGFP (Δ2), or SIRT4(Δ255-314)-eGFP (Δ3) were generated by transfection using the Turbofect reagent (Thermo Fisher Scientific, Oberhausen, Germany). Stable HEK293 cell lines were cultured in selection media containing either G418/Geneticin (400 µg/ml; Genaxxon, Ulm, Germany) or puromycin (1.5 µg/ml; Thermo Fisher Scientific, Oberhausen, Germany). Expression of all SIRT4 constructs was regularly controlled by flow cytometry and/or western blot analysis.

### Preparing total cell lysates for immunoblot analysis

Cells were lysed on ice for 5 min employing a buffer containing 50 mM Tris/HCl (pH 7.4), 100 mM NaCl, 2 mM MgCl_2_, 10% glycerol, 20 mM ß-glycerophosphate, 1 mM Na_3_VO_4_, 1% IGPAL (Thermo Fisher Scientific, Oberhausen, Germany), and 1x protease inhibitor cocktail (Roche). Lysates were cleared by centrifugation (20.000 x g at 4°C for 5 min). Protein concentrations were determined using the Bradford assay.

### Antibodies for immunoblot analysis

Primary antibodies used in western blot analysis include anti-GST (own antibody); anti-GFP (1:1000; #PA1-980, Thermo Fisher Scientific, Oberhausen, Germany); anti-Flag (1:1000; #F742 and #F3165, both from Merck, Darmstadt, Germany); anti-C-RAF-N-terminal (1:1000; #ab181115, Abcam, Cambridge, UK); anti-C-RAF-pS259 (1:1000; #ab173539, Abcam, Cambridge, UK), anti-C-RAF-pY340/341 (1:1000; #sc-16806, Santa Cruz Biotechnology, Heidelberg, Germany); anti-Vinculin (1:1000; #V9131, Merck, Darmstadt, Germany); anti-SIRT4 (1:1000; #69786, Cell Signaling, Frankfurt am Main, Germany); anti-ERK(1/2) (1:1000; #9102, Cell Signaling, Frankfurt am Main, Germany); anti-p-ERK(1/2) (1:1000; #4370, Cell Signaling, Frankfurt am Main, Germany); anti-KRAS (1:1000; 11H35L14, Thermo Fisher Scientific, Oberhausen, Germany). Secondary antibodies employed were from LI-COR (Bad Homburg, Germany; anti-mouse 700 nm: IRDye #926-32213; anti-rabbit 800 nm: IRDye #926-

6807).

### Protein purification

The CRD domain of C-RAF, fused with Glutathione S-transferase (GST), was cloned individually for each single point mutation (A142V, L147F, K148T, I154F, Q156R, L160F, N161Q, E174Q, W187Y), as well as for distinct mutants within set 1 (E174Q, H175R, T178S, K179E, T182L), set 2 (Q156R, F158L, L160F), and set 3 (L147F, K148T, N161Q), using the pGEX-4T1 vector (BioCat GmbH, Heidelberg, Germany). Fusion proteins were expressed in *Escherichia coli* and subsequently purified using glutathione high-capacity magnetic agarose beads (Merck Millipore GmbH, Darmstadt, Germany) following the manufacturer’s guidelines.

### Pull-down assay using GST-fused proteins

Pull-down experiments using GST (Glutathion*-*S*-*Transferase*)-*fused proteins were performed using glutathione agarose beads (Macherey-Nagel, Düren, Germany). The beads were incubated with the GST-fused proteins for 1 h, at 4°C under rotation and centrifuged at 500 g followed by three times washing with ice-cold buffer (30 mM Tris-HCl, 150 mM NaCl, 5 mM MgCl_2_, and 3 mM DTT). In the next step, samples were incubated with total cell lysates from HEK293 cells stably expressing the indicated Flag-tagged sirtuins or SIRT4-eGFP wild-type (WT) and mutants overnight at 4°C followed by three washing steps with ice-cold buffer as indicated above. The protein samples were mixed with Laemmli loading buffer and analyzed by SDS-PAGE and immunoblotting.

### Co-immunoprecipitation analysis

Total cell lysates of HEK293 cells stably expressing C-terminally Flag M2-tagged SIRT4 were incubated overnight at 4°C with anti-Flag M2 agarose beads (Merck, Darmstadt, Germany). The beads were washed three times with washing buffer (50 mM Tris-HCl, 150 mM NaCl, 1 mM EDTA). The beads were mixed with Laemmli loading buffer and co-immunoprecipitation of SIRT4-Flag and endogenous C-RAF proteins was analyzed by SDS-PAGE and immunoblotting. Co-immunoprecipitation of SIRT4-eGFP and endogenous C-RAF using the anti-eGFP nanobody protocol was performed essentially as previously described (33).

### Densitometric analysis of specific immunoblot protein signals followed by statistical evaluation

Intensities of specific protein bands were determined using the Image Studio Lite Version 5.2 software. pull-down data were normalized to the respective total cell lysate signals to ensure an accurate comparison of target protein levels across various samples as previously described (45). Data are presented as mean ± S.D, and One-way ANOVA statistical evaluation was performed using the Origin data analysis software (OriginLab-2021b). Results with at least p ≤ 0.05 were considered significant (*, p ≤ 0.05; **, p ≤ 0.01; and ***, p ≤ 0.001).

### Structural analysis

We created a structural homology model of human SIRT4 on the basis of X-ray diffraction structure of SIRT4 from *Xenopus tropicalis* (PDB ID: 5OJ7) and compared it with human SIRT5 (PDB ID: 4G1C) for the purpose of mutational analysis using PyMOL (Version 4.6.0). Moreover, due to the absence of a complete structure of inactive CRAF, we employed a comparative approach by superimposing the structures of inactive B-RAF (full-length; PDB: 6NYB) to gain insights into the potential structure of inactive C-RAF (46) in complex with 14-3-3. The three-dimensional structure of the resulting inactive state of C-RAF was analyzed and visualized using PyMOL (Version 4.6.0).

### Molecular Docking Simulations

The crystal structures of CRAF_CRD_ (PDB: 1FAQ) and KRAS-CRAF_RBD-CRD_ complex (PDB: 6XHB) were obtained from the Protein Data Bank (PDB) and the human full-length SIRT4(AF-Q9Y6E7) structure was obtained from the AlphaFold database (https://alphafold.ebi.ac.uk/). Molecular docking simulations were performed using default mode settings available in the molecular docking ClusPro 2.0 server (47). From the refined selection of proposed structures, a configuration exhibiting optimal binding energies was chosen, aligning it with experimental data. Subsequent to the docking simulations, the resulting structures were meticulously examined to identify significant molecular interactions using the BIOVIA software.

## Results

### Identification of a selective SIRT4-C-RAF interaction among SIRT and RAF protein family members

In a previous study, we employed mass spectrometry and proteomic analysis to identify novel SIRT4 interacting proteins (33). Interestingly, C-RAF kinase (often referred to by its gene name RAF1), a major component of the MAPK signaling pathway, emerged as a novel SIRT4 binding protein as confirmed by nanobody-based co-immunoprecipitation analysis (**Fig. S1**). Considering the presence of N-terminal regulatory (CR1, CR2) and C-terminal catalytic (CR3) domains in C-RAF (**Fig. 1A**), we hypothesized that the N-terminal CR1 regulatory segment, consisting of the RBD (RAS-binding domain) and CRD (Cysteine-rich domain) domains, might be involved in SIRT4 interaction.

**Figure 1.**
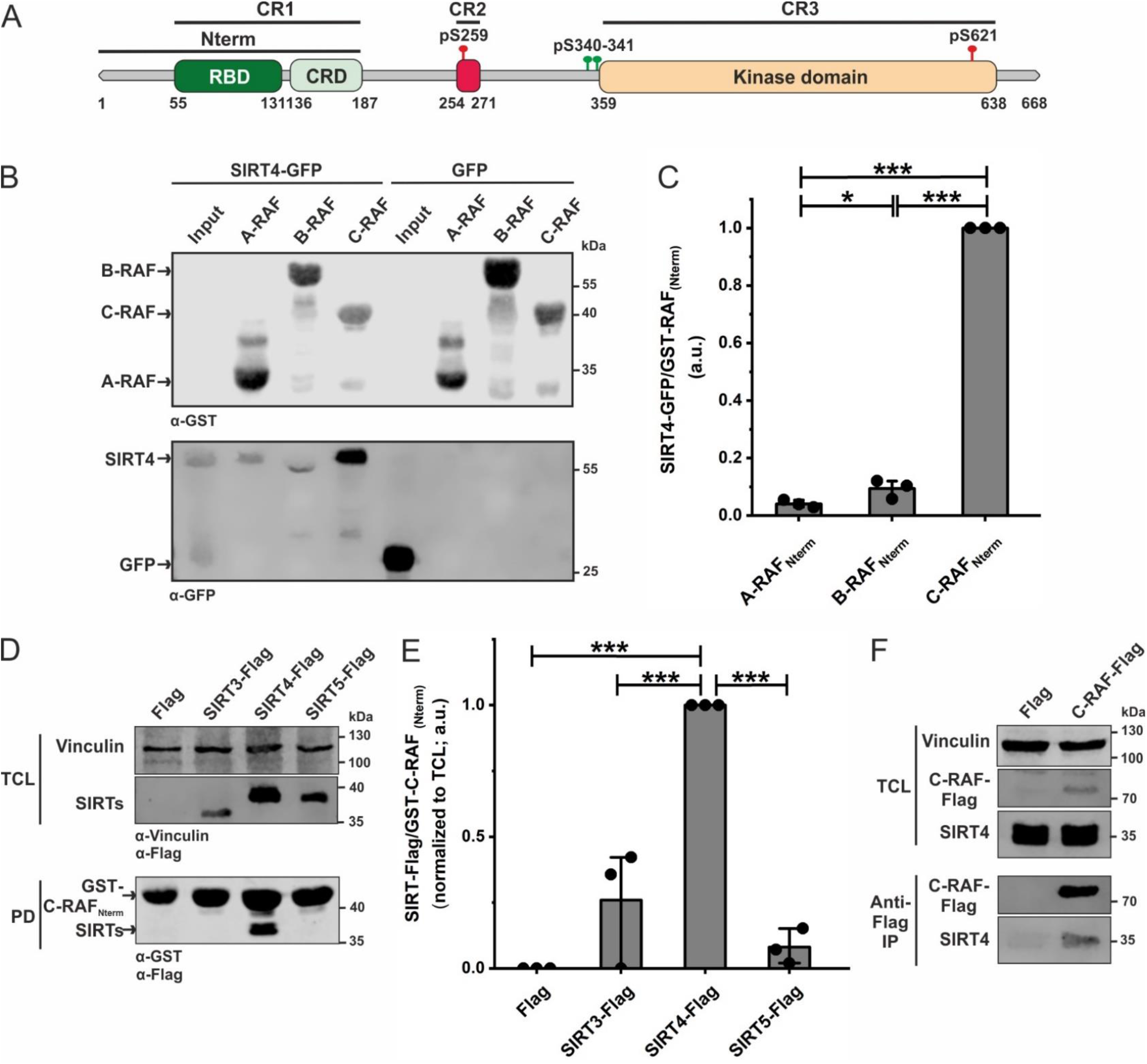
Identification of a selective interaction between SIRT4 and C-RAF within the RAF kinase and SIRT paralogs. (**A**) Domain organization of C-RAF including the RBD (RAS Binding Domain) and CRD (Cysteine-Rich Domain) domains which are parts of the N-terminal region (Nterm). Phosphorylation sites regulating the activity of C-RAF (pS259: inactive form; pY340/341: active form) are indicated. (**B**) Total cell lysates (TCL) from SIRT4-eGFP or eGFP expressing HEK293 cells were subjected to pull-down experiments using normalized bacterial lysates containing the GST-fused Nterm region of A-RAF, B-RAF, or C-RAF. (**C**) Densitometric quantification of immunoblot signals of binding of SIRT4-eGFP to the N-RBD-CRD domain of C-RAF as compared to A-RAF and B-RAF. Data were subjected to statistical One-way ANOVA analysis (mean ± S.D.; *p < 0.05; ***p < 0.001). (**D**) Total cell lysates (TCL) from HEK293 cells expressing Flag-tagged versions of SIRT3, SIRT4, or SIRT5 were subjected to pull-down (PD) experiments using the GST-fused Nterm region of C-RAF. (**E**) Densitometric quantification of immunoblot signals of binding of the Nterm region of C-RAF to SIRT4 as compared to SIRT3 and SIRT5. Data were subjected to statistical One-way ANOVA analysis (mean ± S.D.; ***p < 0.001). (**F**) Co-immunoprecipitation analysis (Anti-Flag Co-IP) of endogenous SIRT4 was performed using total cell lysates (TCL) from Flag-C-RAF expressing COS7 cells.

Accordingly, we addressed the specificity of SIRT4-C-RAF interaction by protein pull-down analysis using bacterially expressed GST-fused N-terminal (Nterm) regions of A-RAF, B-RAF, or C-RAF, each containing the respective RBD and CRD domains. Normalized amounts of GST-RAF lysates were coupled to GSH (glutathione) beads followed by incubation with total cell lysates from HEK293 cells expressing SIRT4-GFP or GFP as control. As indicated in **Fig. 1B, 1C,** and **S2A**, strong physical interaction with SIRT4 was only observed for C-RAF_Nterm_, but not for A-RAF_Ntem_ or B-RAF_Nterm_. In complementary pull-down experiments, we used total cell lysates from HEK293 cells stably expressing C-terminally Flag-tagged SIRT3, SIRT4, or SIRT5. Only SIRT4 exhibited a robust interaction with C-RAF_Nterm_, but not SIRT3 or SIRT5 (**Fig. 1D, 1E, and S2B**). Finally, we immunoprecipitated Flag-tagged C-RAF from COS7 cell lysates and could demonstrate a co-immunoprecipitation of endogenous SIRT4 (**Fig. 1F and S2C**). Overall, our data suggest that within the sirtuin and RAF family members studied, only C-RAF and SIRT4 undergo a unique interaction.

### The CRD domain of C-RAF and the C-terminus of SIRT4 are major determinants of the interaction between SIRT4 and C-RAF

In the next step, we sought to determine the regions or subdomains of C-RAF and SIRT4 that are directly involved in the interaction between these two proteins. We expressed GST-C-RAF-RBD and CRD in *E. coli* and used them to pull down SIRT4-Flag from total cell lysates of HEK293 cells. As indicated in **Fig. S3A-C**, C-RAF_Nterm_ and interestingly CRD alone (C-RAF_CRD_) bound to SIRT4-Flag, although with a higher efficiency seen for C-RAF_Nterm_. However, no or only a slight interaction with SIRT4-Flag could be observed for the RBD (C-RAF_RBD_) (**Fig. 2A, 2B and S3A-C**). These results suggest that the CRD is the major SIRT4-binding domain of C-RAF.

**Figure 2.**
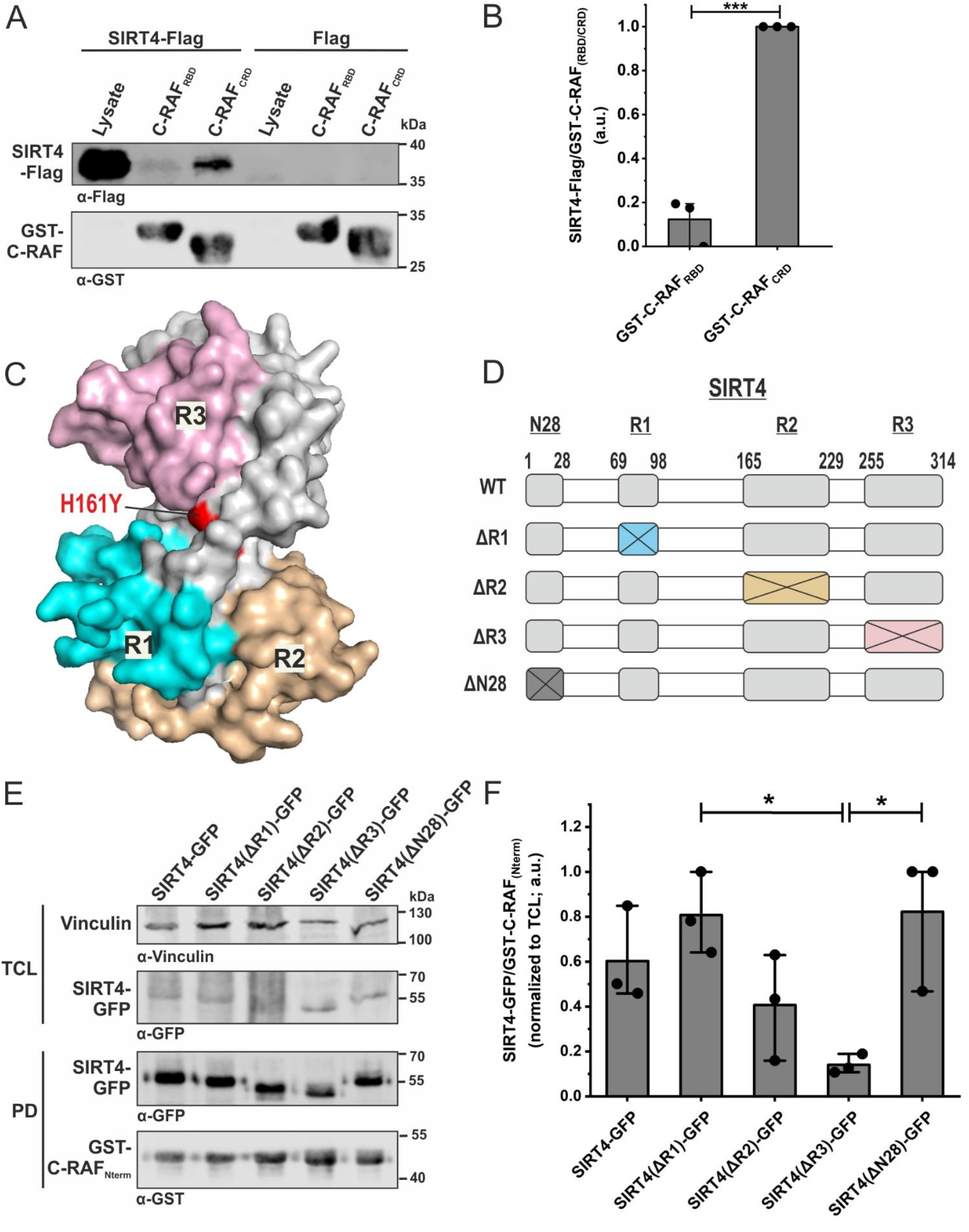
Identification of a selective interaction between the cysteine-rich domain of C-RAF and the very C-terminal region of SIRT4. (**A**) Identification of the CRD domain of C-RAF as the primary SIRT4 interacting domain. Total cell lysates (TCL) from HEK293 cells stably expressing SIRT4-Flag were subjected to pull-down experiments using GST or the GST-fused N-terminal RBD or CRD subdomains of C-RAF. (**B**) Densitometric quantification of immunoblot signals of the relative binding of RBD and CRD subdomains of C-RAF to SIRT4-Flag. Data were subjected to statistical One-way ANOVA analysis (mean ± S.D.; ***p < 0.001). (**C**) The predicted functional surface of SIRT4 was obtained from comparative homology modeling with SIRT5 (**see Fig. S3**). Three regions (R1, R2, and R3), that are different between SIRT4 and SIRT5, are highlighted in the 3D-modelled SIRT4 structure. Replacement of histidine 161 by tyrosine creates the catalytically inactive SIRT4. (**D**) Schematic representation of SIRT4 deletion mutants, including ΔR1, ΔR2, ΔR3, and ΔN28 lacking the N-terminal mitochondrial translocation sequence. (**E**) Equal amounts of total cell lysates (TCL) from HEK293 cells expressing the SIRT4-eGFP of the indicated deletion mutants were subjected to pull-down (PD) analysis using the GST-fused C-RAF_Nterm_. (**F**) Densitometric quantification of immunoblot signals of the relative binding of SIRT4-GFP deletion mutants to the GST-fused C-RAF_Nterm_. Data were subjected to statistical One-way ANOVA analysis (mean ± S.D.; *p < 0.05).

In order to get insight into molecular aspects of SIRT4 binding to C-RAF, we set out to inspect the structures of these proteins and analyze their putative complex. We first generated a homology model of human SIRT4 using the 3D structure of SIRT4 from *X. tropicalis* (PDB: 5OJ7) (48) as a template. Given that SIRT4, but neither SIRT3 nor SIRT5, binds to C-RAF_Nterm_ (**Fig. 1A and 1B**), we have scrutinized their sequences and compared our model structure of SIRT4 with the structure of human SIRT5 (PDB: 4G1C) (**Fig. S4**). This analysis revealed three regions in SIRT4 that differ from SIRT5, i.e., R1(_69-98_), R2(_165-229_), and R3(_255-314_) (**Fig. 2C, 2D**, **and S4**). The corresponding SIRT4 deletion mutants SIRT4(Δ69-98; ΔR1), SIRT4(Δ165-229; ΔR2), and SIRT4(Δ255-314; ΔR3) were generated as C-terminal GFP-tagged proteins, stably expressed in HEK293 cells, and tested for C-RAF_Nterm_ binding in pull-down experiments. As shown in **Fig. 2E, 2F,** and **S3D**, SIRT4(ΔR3) strikingly showed the weakest interaction with C- RAF_Nterm_, whereas ΔR1 and ΔR2 were not significantly different from wild-type SIRT4. Moreover, SIRT4(ΔN28), which lacks the N-terminal mitochondrial translocation signal (44), as well as the catalytically inactive mutant SIRT4(H161Y) (44), bound C-RAF_Nterm_ comparable to wild-type SIRT4 (**Fig. 2E, 2F,** and **S5**). Taken together, C-RAF_CRD_ and the C-terminus of SIRT4, encompassing residues 255-314, are involved in SIRT4-C-RAF interaction, which is independent of the first 28 a.a. of SIRT4 and therefore its mitochondrial localization as well as of the catalytic activity of SIRT4. Our findings also add a new function to the C-terminus of SIRT4 besides its role in proteasomal degradation and stability regulation of SIRT4 (49).

### Mutational analysis of the interaction between C-RAF_CRD_ and SIRT4

We generated nine single mutations and three sets of combined mutations of C-RAF_CRD_ based on the multiple sequence alignment of amino acid deviations of C-RAF_CRD_ in comparison to the CRD domains of A-RAF and B-RAF (**Fig. 3A and 2B**). All mutants were expressed and purified as GST-fusion proteins and subjected to pull-down assays using total cell lysates from SIRT4-Flag expressing HEK293 cells. As indicated in **Fig. 3C and 2E**, and quantitatively analyzed in **Fig. 3D and 3F**, none of the single or combined mutants analyzed decreased interaction of C-RAF_CRD_ with SIRT4-Flag. Rather, we observed significantly stronger binding for the CRD mutants Q156R, E174Q/H175R/T178S/K179E/T182L, and Q156R/F158L/L160F (**Fig. 3C-F**). Thus, these gain-of-function mutations indicate an involvement of the corresponding C-RAF_CRD_ residues in SIRT4 interaction.

**Figure 3.**
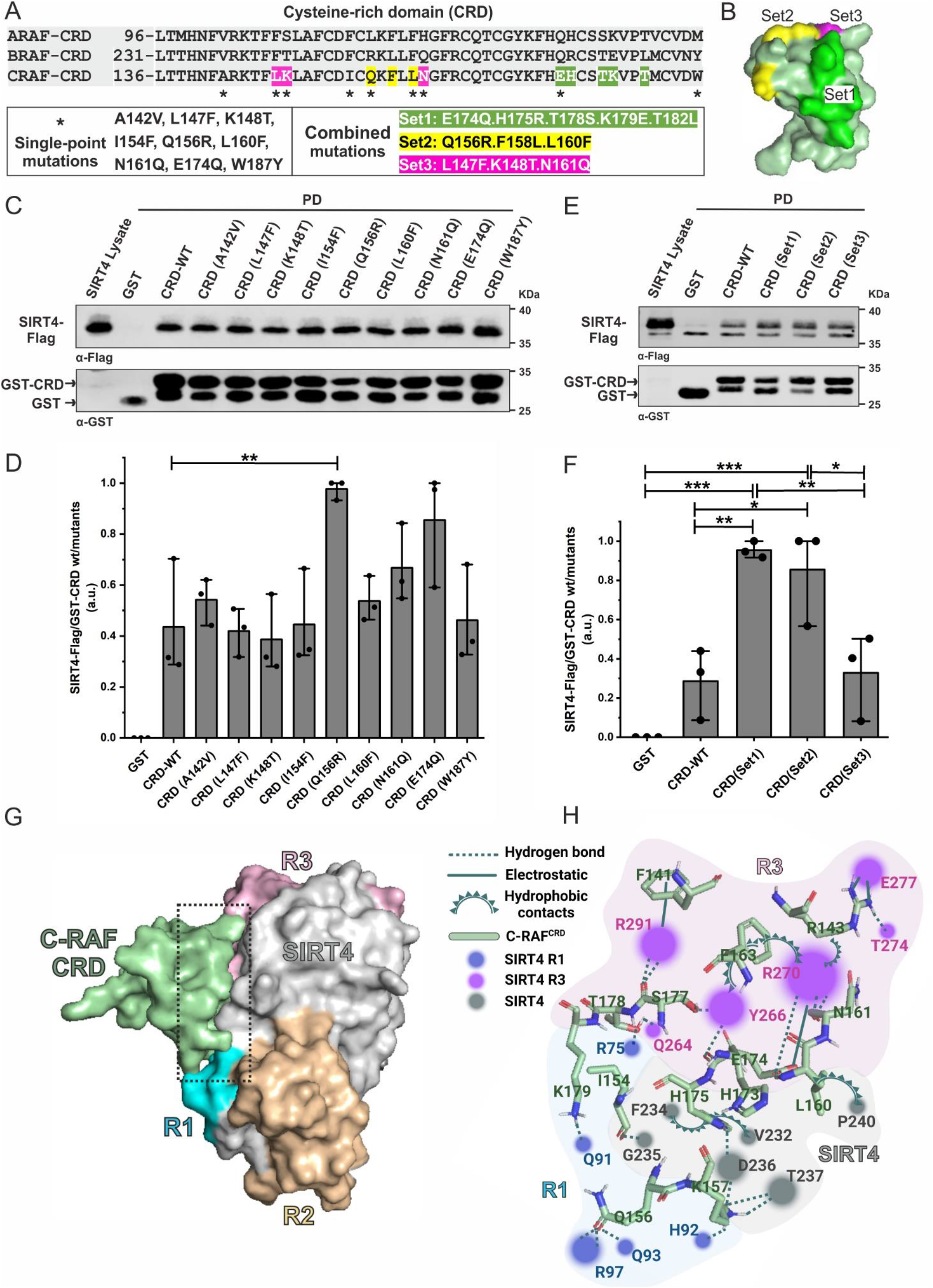
Mapping the SIRT4 binding site of C-RAF. (**A**) Multiple sequence alignment highlights amino acid deviations of the CRD domain of C-RAF as compared to the CRD domains of A-RAF and B-RAF and is the basis for single point and combined mutations of C-RAF generated in this study. (**B**) 3D model of the three sets of combined mutations in the CRD domain of C-RAF. (**C, E**) Total cell lysates (TCL) from HEK293 cells stably expressing SIRT4-Flag were subjected to pull-down experiments using GST, GST-CRD (wildtype), or GST-CRD harboring single point mutations (**C**) or combined mutations (**E**) as indicated in (A). (**D, F**) Densitometric quantification of immunoblot signals of the relative binding of wild-type and mutated CRD subdomains of C-RAF to SIRT4-Flag. Data were subjected to statistical One-way ANOVA analysis (mean ± S.D.; *p < 0.05, **p < 0.01, *** p < 0.001). (**G**) Molecular docking and binding site analysis between the CRD domain of C-RAF and specified regions of SIRT4. The predicted interaction between the CRD domain of C-RAF and the C-terminal region R3 of SIRT4, along with a smaller part of R1, is depicted in this 3D model. (**H**) Schematic, magnified view of the CRD-SIRT4 interacting surface and the involved amino acid residues. The binding types, i.e., hydrogen bonds, electrostatic interactions, and hydrophobic contacts, are indicated. The colored circles mark SIRT4 residues, with the size of the circles indicating the number of interactions with the CRD domain.

### Molecular docking analysis of C-RAF_CRD_ - SIRT4 binding

In order to identify residues of the C-RAF_CRD_ - SIRT4 binding interface and obtain a more detailed insight into their intermolecular interplay, we performed molecular docking analysis between C-RAF_CRD_ (PDB: 1FAQ) and full-length SIRT4 (Q9Y6E7) using the ClusPro 2.0 server. The 3D surface structure (**Fig. 3G**) highlights the binding between C-RAF_CRD_ and R3 of SIRT4, along with certain parts of R1. For a more detailed understanding of this intermolecular binding, analysis of the binding surface utilizing the BIOVIA software revealed an interacting network (**Fig. 3H**), in which the stability of the C-RAF_CRD_ - SIRT4 complex is the result of a combination of various interaction types, *i.e*., hydrogen bonds, electrostatic interactions, and hydrophobic contacts (**Table S1**). For example, the C-RAF_CRD_ residue K157 and the SIRT4 residue D236 form a hydrogen bond with a distance of 1.8 Å, indicative of a strong interaction. C-RAF_CRD_ residues R143, K157, H175, T178, K179, Q156, E174, S177, N161, I154, and SIRT4 residues R75, R97, T274, H92, T237, D236, Q264, Q91, R270, R291, G93, G235 and Y266 further contribute to the binding stability *via* hydrogen bonds. Notably, electrostatic interactions were observed between C-RAF_CRD_ residues R143, E174, and F141 and SIRT4 residues E277, R270, and R291, respectively (**Fig. 3H; Table S1**). Moreover, hydrophobic interactions were identified involving residues of C-RAF_CRD_ (H175, L160, F163, R143) and SIRT4 (V232, F234, P240, Y266, R270).

### SIRT4 binds selectively to the inactive state of C-RAF characterized by phosphorylation of serine 259

C-RAF exists in two distinct forms. Its closed, monomeric, autoinhibited form is stabilized by phosphorylation at serines 259 and 621 (pS259/pS621), and subsequent association with the 14-3-3 dimer (22,50). The C-RAF activation involves a series of complex processes, including dephosphorylation (pS259) and phosphorylation (pY340/pY341) events, conformational changes, dimerization, and association with RAS, 14-3-3, and the membrane, ultimately stabilizing the open, dimeric, active form of C-RAF (15,51–55). Thus, we addressed whether SIRT4 interacts with C-RAF in its active or inactive state. As indicated in **Fig. 4A and S7A**, endogenously expressed C-RAF could be immunoprecipitated from total cell lysates of HEK293 cells expressing SIRT4-Flag. However, when using specific antibodies against pS259-C-RAF (closed, inactive form) and pY340/341-C-RAF (open, active form), only pS259-C-RAF was detected in the immunoprecipitates (**Fig. 4A**). These findings are consistent with homology modeling of C-RAF_CRD_ in the inactive form of C-RAF (**Fig. 4B**), in which the putative SIRT4 binding region remains accessible as part of the CRD domain (indicated in pale green). Moreover, a co-immunoprecipitation of KRAS within the SIRT4-Flag-C-RAF interacting complex could not be detected (**Fig. 4A**), supporting the notion that C-RAF in complex with SIRT4 exclusively exists in its autoinhibited form. Overall, this is consistent with an interaction of KRAS only with the active form of C-RAF, which requires dephosphorylation of S259 and unmasking of the RBD and CRD domains to enable KRAS binding to C-RAF at the membrane [reviewed in (22)]. Further structural analysis offers additional evidence that the SIRT4 binding region of C-RAF_CRD_ contains residues required for KRAS and membrane interaction (**Fig. 4B**).

**Figure 4.**
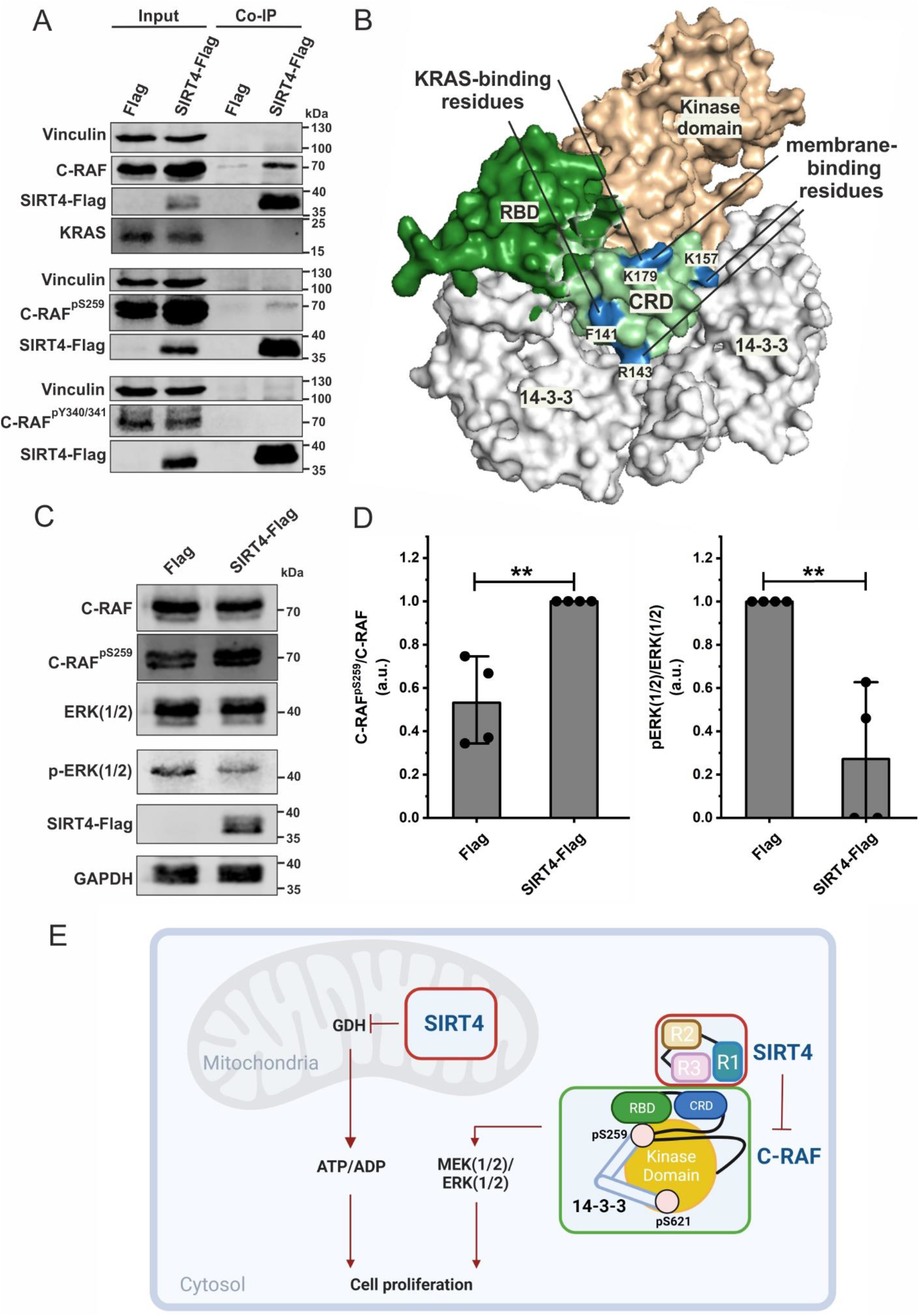
SIRT4 interacts with and upregulates the inactive form of C-RAF phosphorylated at serine 259 (S259). (**A**) Co-immunoprecipitation (Co-IP) analysis using total cell lysates (Input) from HEK293 cells expressing Flag or SIRT4-Flag shows the SIRT4 interaction specifically with C-RAF in its autoinhibited state (pS259-C-RAF) but not with pY340/341-C-RAF in its active state. Moreover, KRAS did not co-immunoprecipitate with the SIRT4-pS259-C-RAF complex. (**B**) A homology model of the closed, inactive C-RAF structure in complex with the 14-3-3 dimer (light grey) was built using the crystal structure of B-RAF as a template. On the left side, the accessibility of the CRD domain in its inactive form is represented (pale green), with highlighted yellow regions indicating the putative binding sites for SIRT4. The model on the right depicts regions highlighted in blue that are crucial for KRAS binding and membrane interaction in the active state of C-RAF. The amino acids involved are indicated. (**C**) Total cell lysates from HEK293 cells expressing Flag or SIRT4-Flag were subjected to immunoblot analysis of pS259-C-RAF and pERK1/2 levels. Ectopic expression of SIRT4 in HEK293 cells increased the levels of inactive pS259-C-RAF and reduced ERK1/2 phosphorylation. (**D**) Densitometric immunoblot analysis of the levels of pS259-C-RAF (left panel) and pERK1/2 (right panel) upon Flag or SIRT4-Flag expression were subjected to statistical One-way ANOVA analysis (mean ± S.D.; **p < 0.01). (**E**) Hypothetical model summarizing the two anti-proliferative axes of SIRT4. SIRT4 displays bifunctional activities in inhibiting glutamate dehydrogenase (GDH) in mitochondria and C-RAF-MAPK signaling in the cytosol. For further explanation, see the discussion.

### SIRT4-C-RAF interaction is associated with the inhibition of the MAPK signaling pathway

It is well established in the literature that SIRT4 overexpression inhibits cell proliferation, among other cellular responses, in various tumor cell lines, most likely through inhibition of the MAPK pathway (37–42). Here, we addressed the regulatory impact of ectopic SIRT4 expression on ERK1/2 phosphorylation. As indicated in **Fig. S7B, 4C and 4D**, ectopic expression of SIRT4 led clearly to an accumulation of the levels of inactive C-RAF phosphorylated at S259. At the same time, MAPK signaling was strongly inhibited as evidenced by an approximately 80% reduction in p-ERK1/2 levels as compared to flag-expressing control cells. Overall, these data indicate that SIRT4 both interacts with and possibly sequesters the inactive form of C-RAF. Thus, the extra-mitochondrial function of SIRT4 towards C-RAF-MAPK signaling may provide a novel control mechanism for tumor suppression (**Fig. 4E**).

## Discussion

The work presented in this study has identified a novel interaction of SIRT4, a tumor suppressor sirtuin, with C-RAF, a key regulatory kinase and a component of the oncogenic MAPK pathway. The results indicate that (i) among the RAF kinases (A-RAF, B-RAF, C-RAF) and sirtuin proteins (SIRT3, SIRT4, SIRT5) analyzed, C-RAF selectively interacts with SIRT4; (ii) this interaction involves the N-terminal CRD domain of C-RAF and in the C-terminal region 3 (R3) of SIRT4 as revealed by pull-down and molecular docking analyses; (iii) mutational analysis of C-RAF_CRD_ identified so far gain-of-function mutations with improved SIRT4 binding, therefore highlighting the importance of these residues in C-RAF_CRD_-SIRT4 interaction; (iv) in particular, SIRT4 specifically interacts with C-RAF in its inactive state (CRAF^pS259^); (v) ectopic expression of functional SIRT4 leads to accumulation of pS259-C-RAF levels, which is associated with inhibition of MAPK signaling as shown by greatly reduced p-ERK1/2 levels. Thus, our data highlight a novel extra-mitochondrial, anti-proliferative function of SIRT4 in binding and potentially sequestering C-RAF from its substrate MEK1/2 and consequently interfering with ERK1/2 activation.

The MAPK signaling pathway plays a crucial role in regulating various cellular processes such as differentiation, survival, and in particular proliferation (56–58). Dysregulation of this pathway is often associated with the development and progression of human diseases, including cancer (4) and developmental disorders like RASopathies (59), the latter exemplified by the RAF1^S257L^ mutation causing cardiomyopathy (60–62). As a key component of the MAPK pathway, C-RAF is activated by upstream receptor-RAS signaling and subsequently activates various downstream effectors, particularly MEK1/2 kinases and thus ERK1/2 signaling (21,22,57). Several studies have highlighted the molecular mechanism of C-RAF regulation underlying post-translational modifications by phosphorylation and dephosphorylation, auto-inhibition, and conformational changes accompanied with stabilized protein-protein interaction (12,17–19). Classically, RAS proteins and 14-3-3 binding are major regulators of RAF activation, membrane recruitment of C-RAF, and its stability (8,22). The complexity of C-RAF regulation is further highlighted by its heterodimerization with B-RAF which acts as an allosteric inducer of C-RAF in normal and cancer cells in a RAS-independent manner (63).

Recent findings have identified and elaborated further C-RAF regulators. SHOC2 serves as a scaffold protein for C-RAF, which recruits together with MRAS protein phosphatase 1 to dephosphorylate inactive C-RAF at S259, thereby facilitating C-RAF interaction with RAS at the plasma membrane (64–66). In another example, SHOC2 serves as a regulatory factor for C-RAF and has been shown to accelerate the interaction between RAS and C-RAF, ultimately influencing the spatiotemporal patterns of the RAS-ERK signaling pathway (66). Moreover, RKTG (RAF kinase trapping to Golgi) has been suggested to regulate the spatial localization of C-RAF by trapping it to the Golgi, thereby altering the interaction of C-RAF with RAS and MEK1 and inhibiting ERK signaling (67). Another regulator of C-RAF is RKIP (25–28) which binds to the N-terminal region of C-RAF and thereby inhibits C-RAF-mediated phosphorylation and activation of MEK1/2 (29,68). Interestingly, a comparison between RKIP and SIRT4 reveals cellular and functional similarities: (i) both proteins are tumor suppressors (28,69) inhibiting/preventing C-RAF activation, and their expression is usually down-regulated in cancer (27,37–39), although the underlying mechanisms for SIRT4 are yet unclear; (ii) SIRT4 and RKIP are both involved in the regulation of mitotic cell division. SIRT4 achieves this through centrosomal localization and control of microtubule dynamics (33), while RKIP achieves this by interacting with aurora-B and control of the mitotic checkpoint (70); lastly, (iii) both SIRT4 (44,71) and RKIP are linked to the regulation of autophagy. RKIP is involved in LC3 processing and presumably contributes to autophagosome formation upon starvation (72,73). The role of the SIRT4-C-RAF axis in regulating these cellular responses requires further characterization.

The intermolecular interplay within the C-RAF_CRD_ - SIRT4 binding interface still needs to be determined at the residual level. The single and combined C-RAF_CRD_ mutations, that were defined based on homology comparison to the CRD domains of A-RAF and B-RAF (which do not interact with SIRT4), did not interfere with the C-RAF_CRD_-SIRT4 interaction (**Fig. 3**). Therefore, molecular docking experiments of C-RAF_CRD_ on SIRT4 were performed to determine their putative binding interface. These analyses revealed besides residues from the mutational analysis of the C-RAF_CRD_ domain (**Fig. 3**) further candidate residues that may be critical for SIRT4 interaction (**Fig. 3H** and **Table S1**). In addition, candidate residues within the R3 and R1 regions of SIRT4 could be identified, whose function also needs to be tested by mutational analysis.

Interestingly, the SIRT4 binding region of C-RAF_CRD_ comprises residues which are also required for KRAS and membrane interaction of C-RAF_CRD_ (**Fig. 4B**). Previous results identified seven essential basic residues within the CRD domain (R143, K144, K148, K157, R164, K171, and K179) which are pivotal for membrane interaction, with particular emphasis on the key basic residues R143, K144, and K148 (24). R143, K157, and K179 are accessible in the inactive state of C-RAF and part of the SIRT4 interaction surface, while the remaining residues are located on the opposite side and are shielded by 14-3-3 dimers (**Fig. 4B**). In terms of KRAS binding, F141 and K179 are vital for the interaction of KRAS and C-RAF during the activation process (8). In the inactive state of C-RAF, in addition to K179, F141 is also accessible in the CRD domain, consistent with the involvement of these two residues in SIRT4 binding as revealed by docking analysis (**Fig. 4B**).

At the level of the functional C-RAF-SIRT4 interplay, it is currently unknown whether C-RAF is regulated by an acetylation/deacetylation cycle and whether C-RAF represents an enzymatic target of SIRT4. SIRT4 itself displays different NAD^+^-dependent enzymatic activities, including ADP-ribosylation, diacylation, and deacetylation (74), with recent findings indeed uncovering several new SIRT4 deacetylation targets not only inside, but also outside of mitochondria (35,75). In this regard, there is a paradigm for the regulation of B-RAF by SIRT1. Acetylation of B-RAF at lysine 601 by the p300 acetyltransferase promotes B-RAF kinase activity, thereby enhancing the proliferation melanoma and resistance to B-RAF^V600E^ inhibitors (76). On the contrary, SIRT1 deacetylates B-RAF at K601 and thereby inhibits proliferation. Thus, SIRT1 mediates hypo-acetylation of B-RAF and thereby (fine) regulates its downstream MAPK signaling activity.

Our findings introduce another layer of complexity to the regulatory network of C-RAF and MAPK signaling by identifying SIRT4 as a C-RAF binder specifically in its inactive state. As summarized in **Fig. 4E**, in mitochondria, SIRT4 inhibits anaplerosis and ultimately ATP generation *via* inhibition of GDH (31). Outside of the mitochondria, SIRT4 interacts, seemingly *via* its C-terminal R3, with the inactive ‘closed’ form of C-RAF, in which the kinase domain is concealed through 14-3-3 binding to pS259 and pS621. SIRT4 binding to the CRD domain of C-RAF potentially stabilizes pS259/pS621-C-RAF thereby preventing membrane recruitment, which is followed by RAS binding, and activation of C-RAF. Consequently, association of SIRT4 with C-RAF interferes with the activation of downstream MEK/ERK signaling, consistent with findings showing SIRT4 mediated inhibition of the MAPK pathway (37–42).

So far, only the MEK1/2 kinases have been well-described as substrates of C-RAF. However, there are several indications of kinase-independent functions/activities of C-RAF, including regulation of apoptosis, cell cycle progression, and migration (77). In this context, there is a broad spectrum of C-RAF targets that could interact either directly or indirectly with active (pSer-338) or inactive (pSer-259) forms of C-RAF. This interaction could also be RAS-dependent or independent. For example, the interaction between MST2 and C-RAF (pSer-259) prevents MST2 dimerization (12) and consequently modulates the strength of mitotic and apoptosis signaling. Of note, we also observed an impact of ectopic SIRT4 expression on the Hippo tumor suppressor pathway that besides the MAPK pathway also regulates cell proliferation (78,79). In particular, the increase of pS259-C-RAF levels upon SIRT4 expression (**Fig. 4C** and **4D**) was associated with a decrease in the pYAP/YAP ratio (unpublished results). Taken together, we describe a novel SIRT4-CRAF axis that negatively impacts both MAPK and Hippo-YAP signaling. Another example is ASK1, which normally activates the pro-apoptotic signaling pathways JNK and p38, and is negatively regulated by C-RAF (80). C-RAF phosphorylated at residue 338 interacts with the N-terminal domain of ASK1 in a kinase-independent and HRAS-dependent manner (81)). The C-RAF-ASK1 complex formed in mitochondria is disrupted by oxidative stress (82). Whether SIRT4 plays a role in this process remains to be investigated. Further C-RAF activities that may be affected by SIRT4 include the phosphorylation of the PKCθ-BAD complex in the control of anti-apoptosis responses (83), stimulation of negative regulation of cell migration through direct interaction with ROCKα (84), promotion of the cell cycle progression through interaction with Polo-Like kinase 1 (PLK1) and Aurora kinase A at the mitotic spindle, and the regulation of the DNA damage response through interaction with Checkpoint kinase 2 (CHK2) (85–88).

## Author Contributions Statement

M.R.A. and R.P.P. initiated the project and designed the study. M.M., A.L., P.V., M.R.A., and R.P.P. designed, performed, and analyzed the experiments. M.M., R.D., and S.N-R. performed the homology modeling and molecular docking simulation analyses. M.M., M.R.A., and R.P.P. wrote the paper. All authors read, discussed, corrected, and approved the final version of the manuscript.

## Acknowledgments

We are grateful to our colleagues from the Institute of Biochemistry and Molecular Biology II for their support, helpful advice, and fruitful discussions. We thank Dr. Motta (Genetics and Rare Diseases Research Division, Rome) for providing the Flag-C-RAF expression vector. This study was supported by the German Research Foundation (DFG; grant AH 92/8-3 to M.R.A.), the European Network on Noonan Syndrome and Related Disorders (NSEuroNet, grant 01GM1602B to M.R.A.), and in part by the Foundation for Ageing Research of the Heinrich Heine University (grants 701.810.783 to R.P.P., and 701.810.845 to M.R.A.).

## Supplementary information

**Table S1:** Molecular docking analysis of C-RAF_CRD_-SIRT4 binding and summary of interacting amino acid residues and binding types. The amino acids highlighted in gray belong to SIRT4 (excluding R1/2/3), the blue residues represent R1, and the pink residues represent R3 within the binding site.

| SIRT4 | C-RAF<sup>CRD</sup> | Category | Distance [Å] |
| --- | --- | --- | --- |
| V232 | H175<sup>Set1</sup> | Hydrophobic | 4.78902 |
| F234 | H175<sup>Set1</sup> | Hydrophobic | 4.58657 |
| P240 | L160<sup>*/Set2</sup> | Hydrophobic | 5.31559 |
| G235 | I154<sup>*</sup> | Hydrogen Bond | 3.51669 |
| T237 | K157 | Hydrogen Bond | 2.86838 |
| T237 | K157 | Hydrogen Bond | 2.84552 |
| D236 | H175<sup>Set1</sup> | Hydrogen Bond | 2.05588 |
| D236 | K157 | Hydrogen Bond; Electrostatic | 1.79794 |
| H92 | K157 | Hydrogen Bond | 1.90453 |
| Q91 | K179<sup>Set1</sup> | Hydrogen Bond | 1.65633 |
| G93 | Q156<sup>*/Set2</sup> | Hydrogen Bond | 3.51031 |
| R97 | Q156<sup>*/Set2</sup> | Hydrogen Bond | 1.73351 |
| R97 | Q156<sup>*/Set2</sup> | Hydrogen Bond | 2.0763 |
| R75 | T178<sup>Set1</sup> | Hydrogen Bond | 1.73089 |
| Y266 | F163 | Hydrophobic | 4.08338 |
| R270 | R143 | Hydrophobic | 5.00044 |
| R270 | F163 | Hydrophobic | 5.18852 |
| R270 | N161<sup>*/Set3</sup> | Hydrogen Bond | 2.40989 |
| R270 | N161<sup>*/Set3</sup> | Hydrogen Bond | 2.7401 |
| Q264 | T178<sup>Set1</sup> | Hydrogen Bond | 2.00436 |
| Y266 | E174<sup>*/Set1</sup> | Hydrogen Bond | 2.72696 |
| Y266 | S177 | Hydrogen Bond | 2.02702 |
| T274 | R143 | Hydrogen Bond | 1.72576 |
| R291 | S177 | Hydrogen Bond | 1.85345 |
| R291 | S177 | Hydrogen Bond | 2.94189 |
| R270 | E174<sup>*/Set1</sup> | Hydrogen Bond; electrostatic | 1.81366 |
| R270 | E174<sup>*/Set1</sup> | Electrostatic | 2.72195 |
| E277 | R143 | Electrostatic | 2.85759 |
| E277 | R143 | Electrostatic | 2.74066 |
| R291 | F141 | Electrostatic | 4.73388 |

\*: Residues considered for single-point mutations
Set1/2/3: specific residues targeted for combined mutations

**Figure S1.**
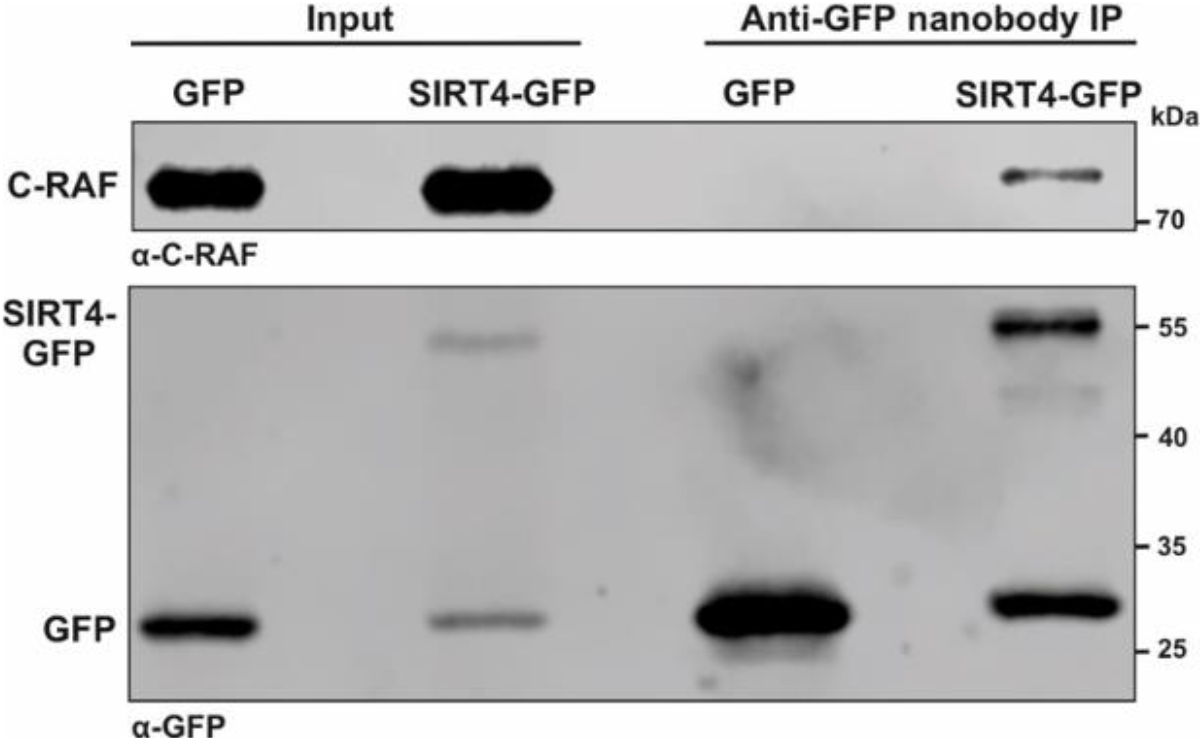
Validation of C-RAF as SIRT4 interacting protein. Total cell lysates of HEK293 cells stably expressing SIRT4-GFP or GFP as control were subjected to co-immunoprecipitation (IP) analysis using the anti-GFP nanobody method followed by immunoblotting for endogenous C-RAF using an Abcam antibody (#ab181115).

**Figure S2.**
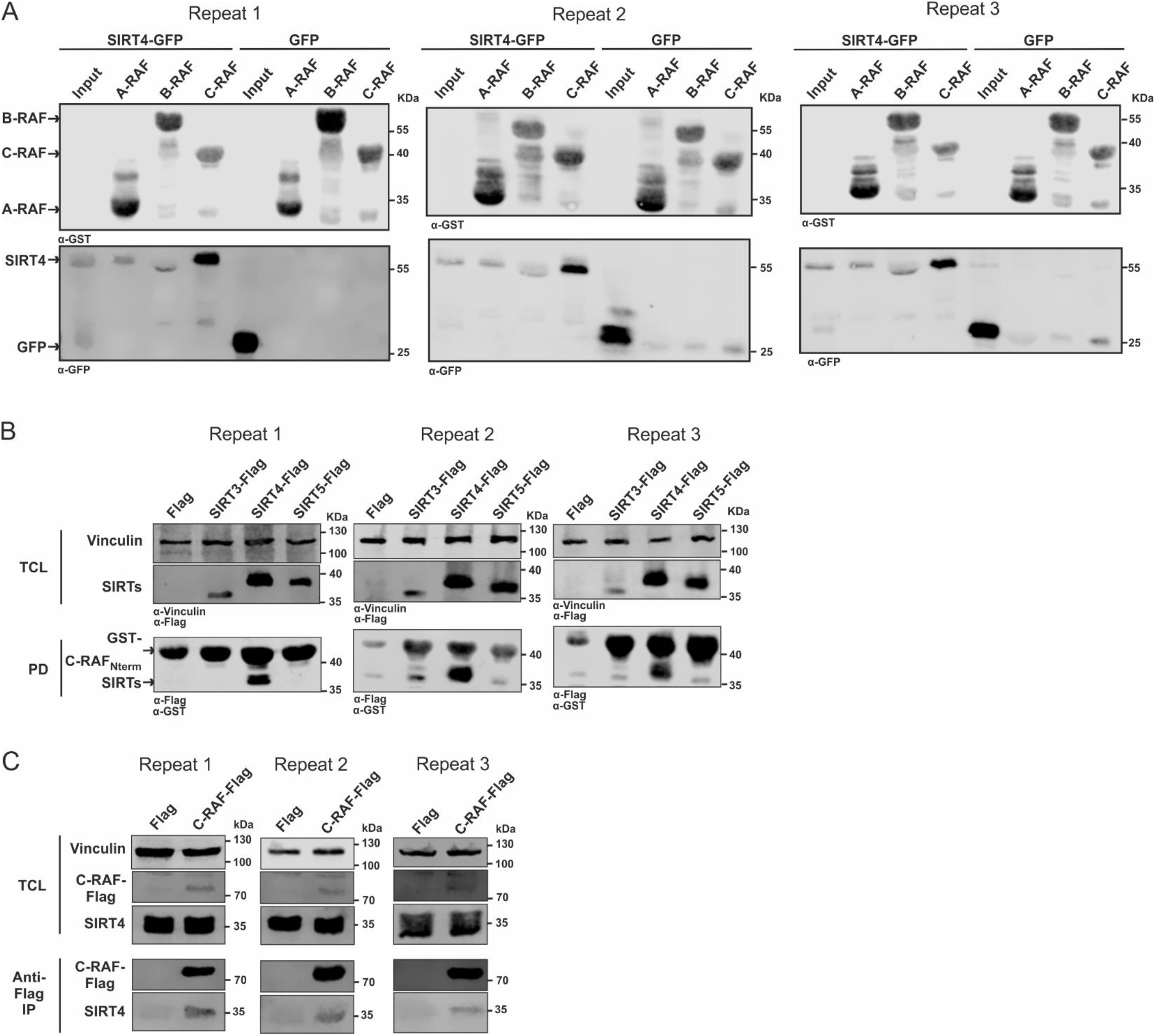
The independent experimental repeats statistically analyzed in **Fig. 1B-F** are depicted. (**A**) Single experiments related to **Fig. 1B, C**. (**B**) Single experiments related to **Fig. 1D,E**. (**C**) Single experiments related to Fig. 1F.

**Figure S3.**
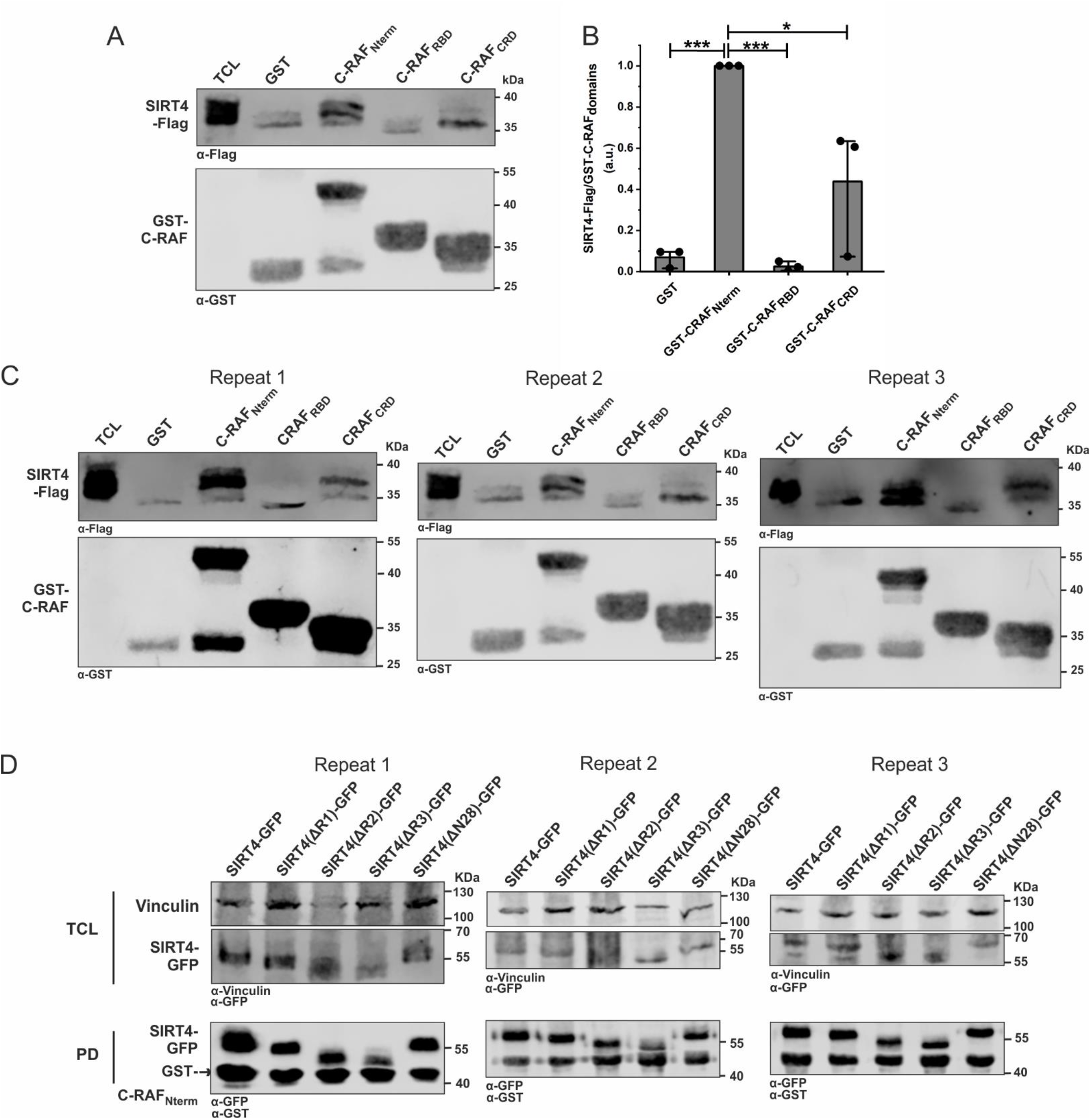
The CRD domain within the N-terminus of C-RAF interacts with SIRT4. (**A**) Total cell lysates (TCL) from HEK293 cells stably expressing SIRT4-Flag were subjected to pull-down experiments using GST, GST-C-RAF_Nterm_ or GST-C-RAF_CRD_. (**B**) Densitometric quantification of immunoblot signals of binding of N-RBD-CRD subdomains of C-RAF to SIRT4-Flag. Data were subjected to statistical One-way ANOVA analysis (mean ± S.D.; *p < 0.05; ***p < 0.001). (**C**) The three independent experimental repeats statistically analyzed in **Fig. S2A,B** are depicted. (**D**) The three independent experimental repeats statistically analyzed in **Fig. 2E,F** are depicted.

**Figure S4.**
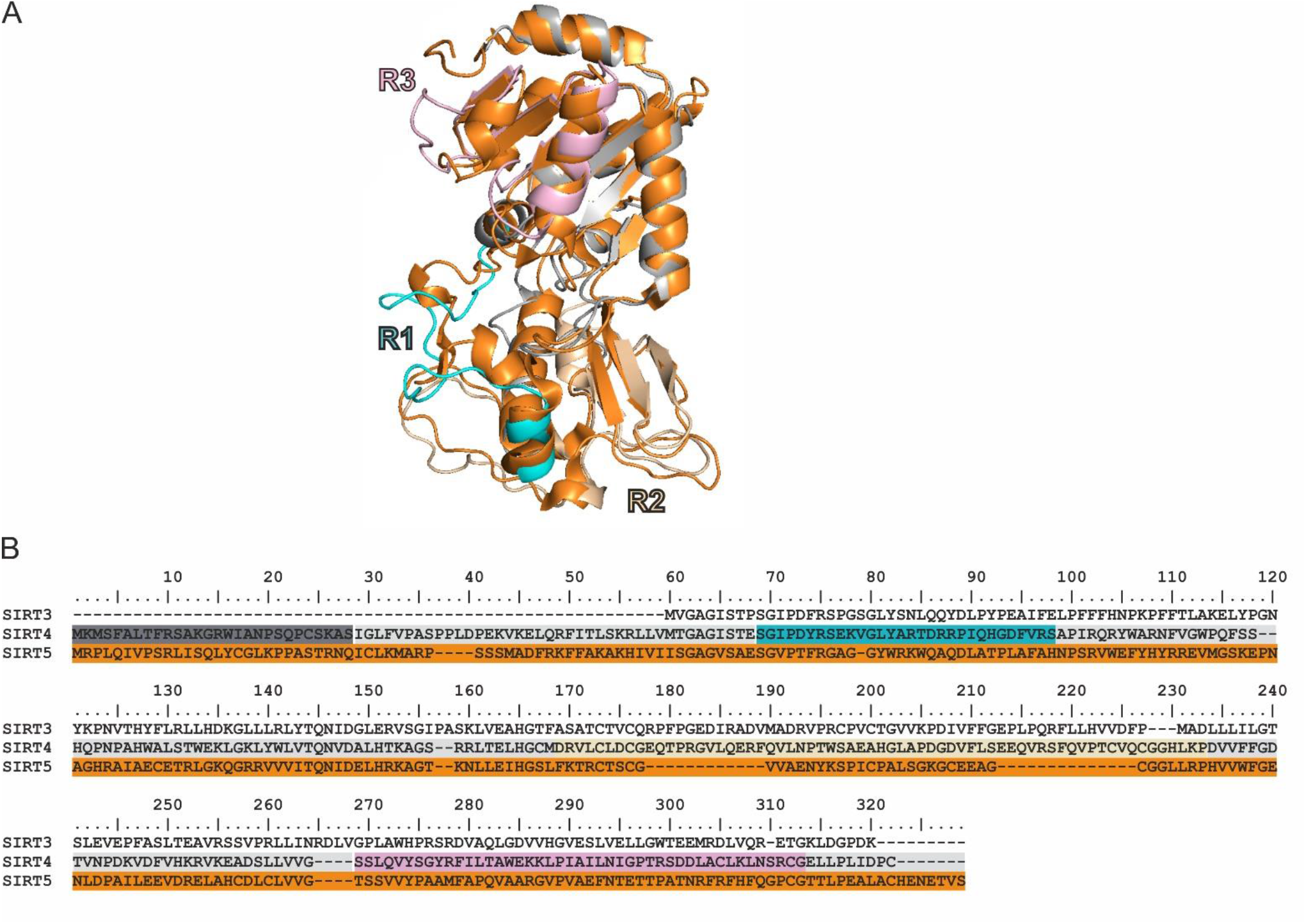
Homology modeling of human SIRT4 and SIRT5 proteins. (**A**) 3D homology comparison between SIRT5 (orange mesh) and SIRT4 (light gray mesh). The SIRT4 regions are highlighted in colors corresponding to **B**. (**B**) Sequence alignment of human SIRT3, SIRT4, and SIRT5. The alignment highlights the initial N-terminal 28 amino acids of SIRT4 in gray. These amino acids are not represented in the structure. Additionally, regions R1, R2, and R3 are indicated by cyan, mustard, and pink frames, respectively.

**Figure S5.**
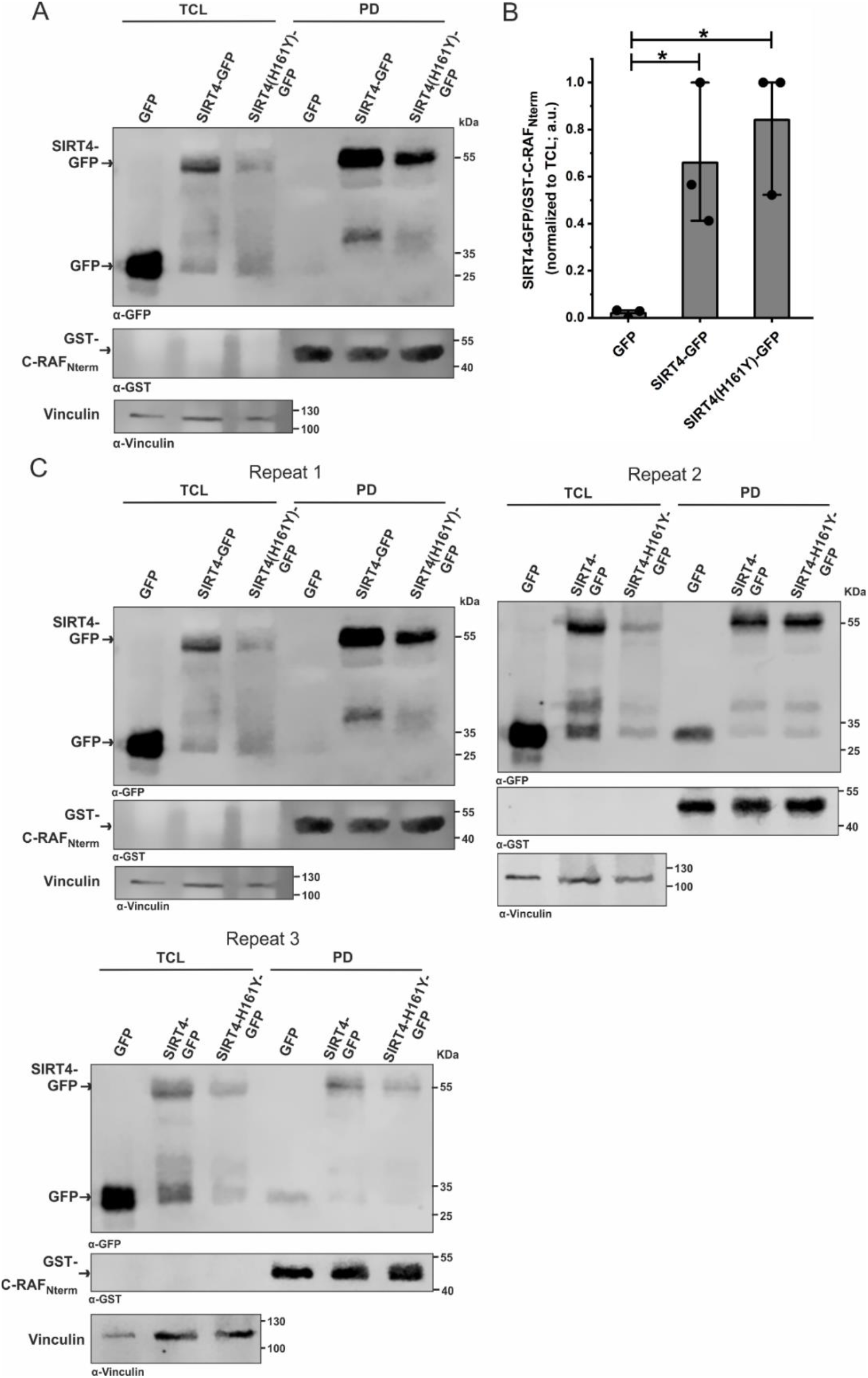
The interaction between SIRT4 and C-RAF is independent of the catalytic activity of SIRT4. (**A**) Total cell lysates (TCL) from HEK293 cells expressing GFP, SIRT4-GFP, or the catalytically inactive mutant SIRT4(H161Y)-GFP were subjected to pull-down (PD) experiments using the GST-fused Nterm domain of C-RAF. (**B**) Densitometric quantification of immunoblot signals of binding of the Nterm domain of C-RAF to GFP, SIRT4-GFP, or SIRT4(H161Y)-GFP. Data were subjected to statistical One-way ANOVA analysis (mean ± S.D.; *p < 0.05). (**C**) The three independent experimental repeats statistically analyzed in (**B**) are depicted.

**Figure S6.**
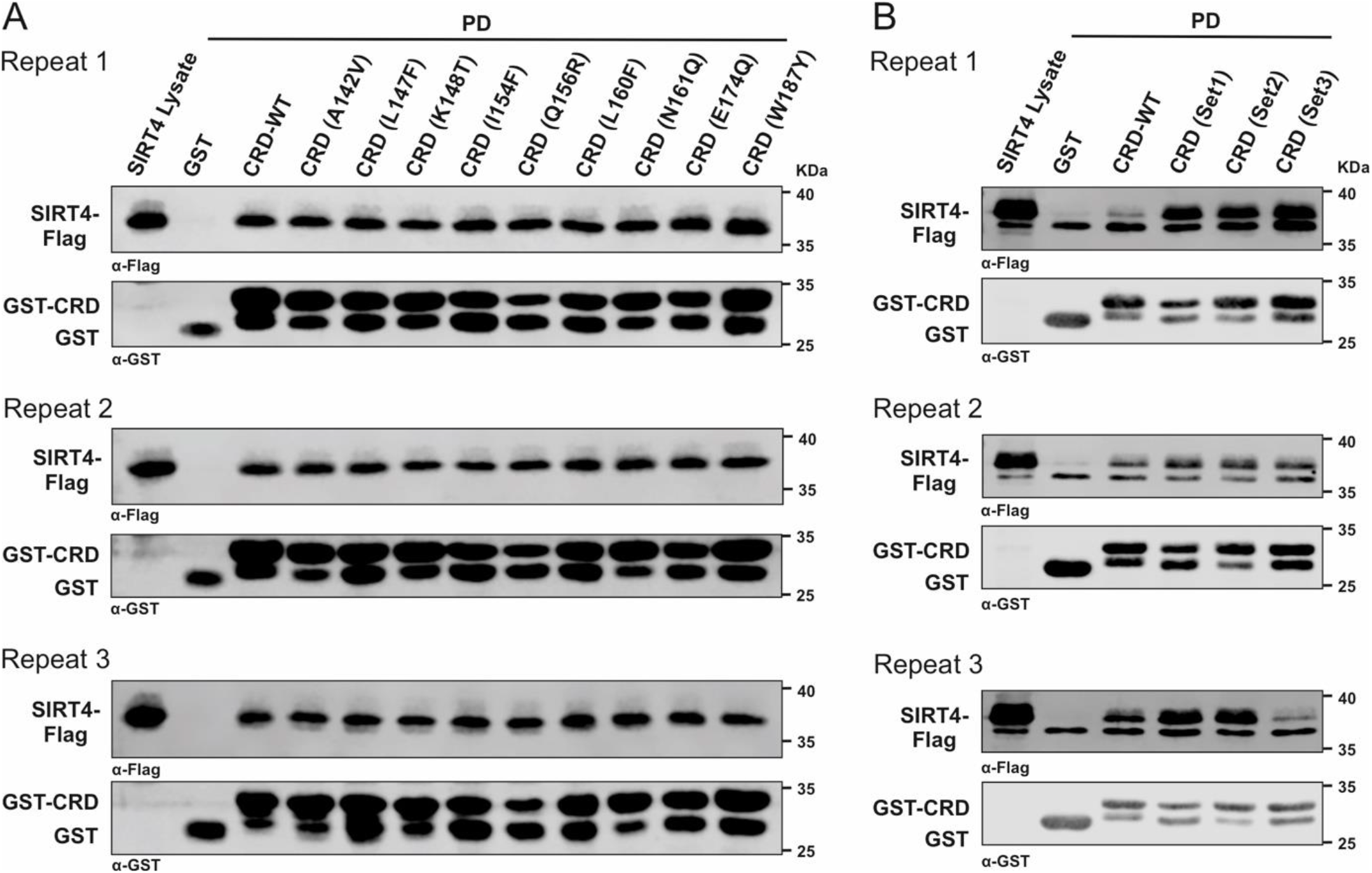
The three independent experimental repeats statistically analysed in **Fig. 3** are depicted. (**A**) analysis of CRD single point mutants (**Fig. 3C,D**). (**B**) analysis of CRD forms with combined mutations (**Fig. 3E,F**).

**Figure S7.**
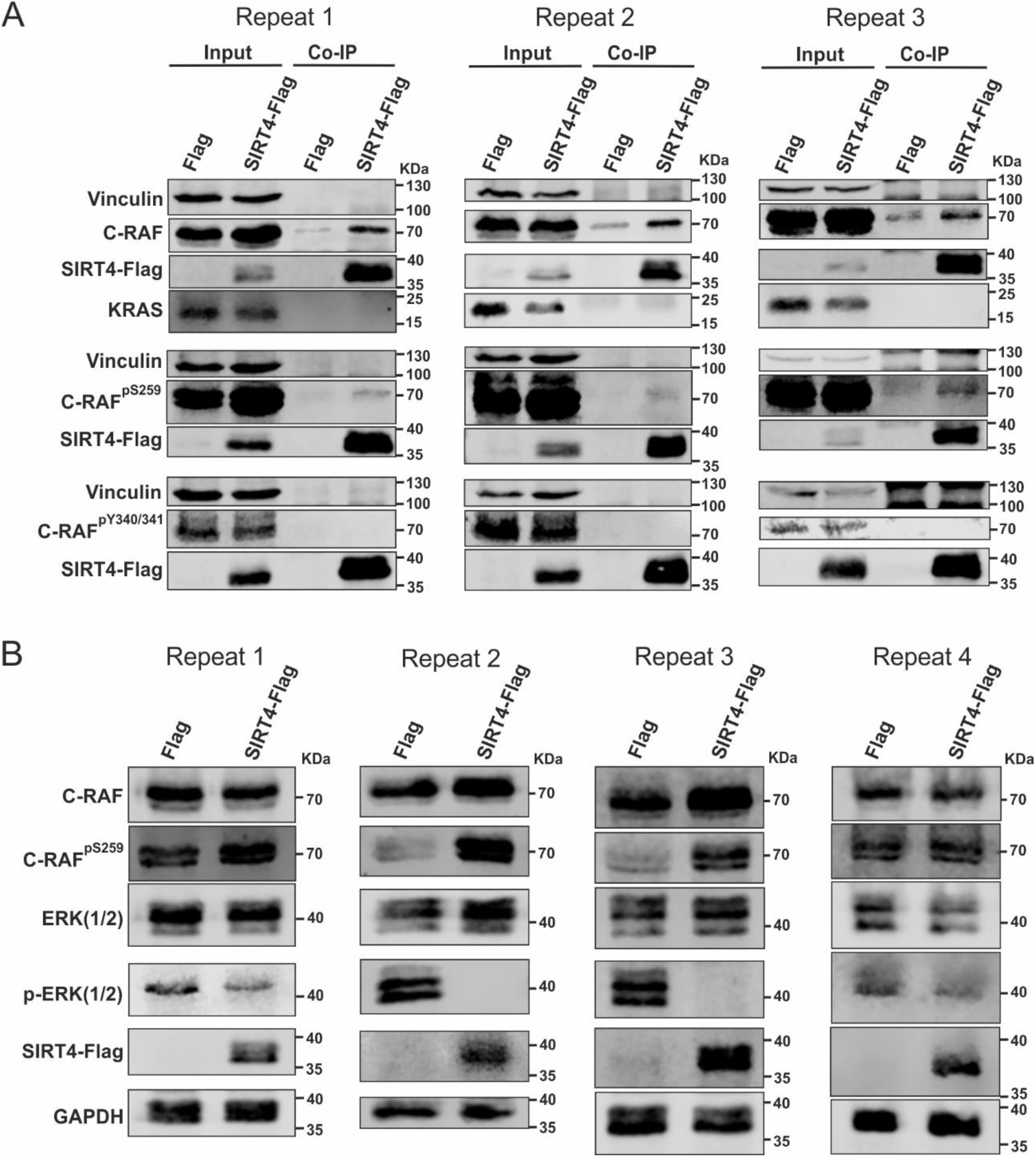
The experimental repeats statistically analyzed in **Fig. 4** are depicted. (**A**) Interaction of SIRT4 with the inactive form of C-RAF phosphorylated at serine 259 (S259) (Fig. 4A). (**B**) SIRT4 expression upregulates protein levels of C-RAF phosphorylated at serine 259 (S259) and downregulates pERK1/2 levels (**Fig. 4C**).

